# miR-146a-5p mediates inflammation-induced β cell mitochondrial dysfunction and apoptosis

**DOI:** 10.1101/2024.03.18.585543

**Authors:** Preethi Krishnan, Renato Chaves Souto Branco, Staci A. Weaver, Garrick Chang, Chih-Chun Lee, Farooq Syed, Carmella Evans-Molina

## Abstract

We previously showed that miR-146a-5p is upregulated in pancreatic islets treated with pro-inflammatory cytokines. Others have reported that miR-146a-5p overexpression is associated with β cell apoptosis and impaired insulin secretion. However, the molecular mechanisms mediating these effects remain elusive. To investigate the role of miR-146a-5p in β cell function, we developed stable MIN6 cell lines to either overexpress or inhibit the expression of miR-146a-5p. Monoclonal cell populations were treated with pro-inflammatory cytokines (IL-1β, IFN*γ*, and TNFα) to model type 1 diabetes (T1D) *in vitro*. We found that overexpression of miR-146a-5p increased cell death under conditions of inflammatory stress and led to mitochondrial membrane depolarization, whereas inhibition of miR-146a-5p reversed these effects. Additionally, inhibition of miR-146a-5p increased insulin secretion, mitochondrial DNA copy number, respiration rate, and ATP production Further, RNA sequencing data showed enrichment of pathways related to insulin secretion, apoptosis, and mitochondrial function when the expression levels of miR-146a-5p were altered. Finally, a temporal increase in miR-146a-5p expression levels and a decrease in mitochondria function markers was observed in islets derived from NOD mice. Collectively, these data suggest that miR-146a-5p may promote β cell dysfunction and death during inflammatory stress by suppressing mitochondrial function.

## INTRODUCTION

Type 1 diabetes (T1D) is characterized by immune-mediated destruction of pancreatic β cells and manifests clinically after a substantial loss of β cell mass and function, leading to severe hyperglycemia and metabolic instability (1). Evidence from clinical studies suggests that β cell loss may begin years before clinical diagnosis, and disease initiation is documented by the development of autoantibodies against β cell antigens, such as GAD, insulin, IA-2, and ZnT8 (2). Evidence from birth cohort studies shows that 85% of individuals positive for two or more autoantibodies will develop T1D within 15 years of follow-up (3).

Immune-mediated destruction of β cells in T1D is accompanied by the activation of a series of transcriptional and signaling pathways which are thought to render the β cell more immunogenic while also invoking self-imposed dysfunction and/or death. Inflammatory insults such as pro-inflammatory cytokines contribute to T1D pathogenesis, and others have shown that treatment of mouse and human pancreatic islets and β cell lines with cytokines (IL1β, IFN*γ*, TNFα, IFNα) recapitulates many of the *in vivo* molecular changes seen in early and late stages of T1D (4, 5).

In addition to alterations in protein-coding genes, non-protein-coding genes, particularly microRNAs (miRNAs), contribute to T1D development (6–8). miRNAs are small non-coding RNAs that serve as global regulators of gene expression (9–11). miRNAs have been implicated in a variety of biological processes within the β cell, including endocrine cell specification (12), β cell development (13), insulin secretion (14), glucose regulation (15), and apoptosis (16). Previously, we identified miR-146a-5p as a cytokine-modulated miRNA in human islets treated with pro-inflammatory cytokines. miR-146a-5p was coordinately upregulated in human islets and islet-derived extracellular vesicles (6, 7). Notably, we observed an increase in expression of miR-146a-5p in plasma-derived extracellular vesicles from individuals with autoantibody-positivity and normoglycemia and in recent onset T1D, suggesting that miR-146a-5p may have utility as a T1D biomarker (7).

Previous research has demonstrated that robust and sustained upregulation of miR-146a-5p during inflammatory states led to an increase in apoptosis and a significant decrease in insulin secretion in MIN6 cells (17). However, the mechanisms through which miR-146a-5p exert these effects remain elusive. Data from other inflammatory disease models classify miR-146a-5p as a mitomiR—a miRNA that is enriched in the mitochondria, where it is implicated to play a critical role in the regulation of mitochondrial function (18–20). Moreover, inflammation-mediated mitochondrial dysfunction has been established as a pivotal factor in the pathogenesis of T1D. (21–23). Based on this premise, our study aims to investigate whether miR-146a-5p contributes to T1D pathogenesis through dysregulation of mitochondrial function.

## RESULTS

### miR-146a-5p overexpression promotes β cell death under pro-inflammatory conditions

Our previous RNA sequencing analysis of human islets exposed to pro-inflammatory cytokines revealed a significant upregulation of miR-146a-5p during inflammation (7). In agreement with these results, mouse islets (Fig. 1A) and MIN6 cells (Fig. 1B) that were exposed to pro-inflammatory cytokines (IL1β, IFN*γ*, and TNFα) for 24 hours showed a significant increase in the expression of miR-146a-5p. Next, to functionally delineate the role of miR-146a-5p in MIN6 cells, we established stable cell lines using a lentiviral vector. These included a lentiviral vector expressing a scrambled sequence with no similarity to any known mouse genomic sequence (Scr), a lentiviral vector overexpressing miR-146a-5p (OE), and a lentiviral vector with anti-sense miR-146a-5p (ZIP). qRT-PCR analysis of OE cells showed a significant upregulation of miR-146a-5p, while miR-146a-5p expression levels remained unchanged in cells transduced with the ZIP knockdown vector compared to Scr control (Fig. 1C). Furthermore, there was an approximately 500-fold increase in miR-146a-5p levels in Scr cells treated with cytokines compared to untreated Scr cells (Fig. S1A). In contrast, we only observed a 2-fold increase in miR-146a-5p levels in ZIP cells treated with cytokines compared to untreated ZIP cells (Fig. S1B). Likewise, we observed an approximately 8-fold increase in miR-146a-5p levels in OE cells treated with cytokines compared to untreated OE cells (Fig. S1C).

**Figure 1.**
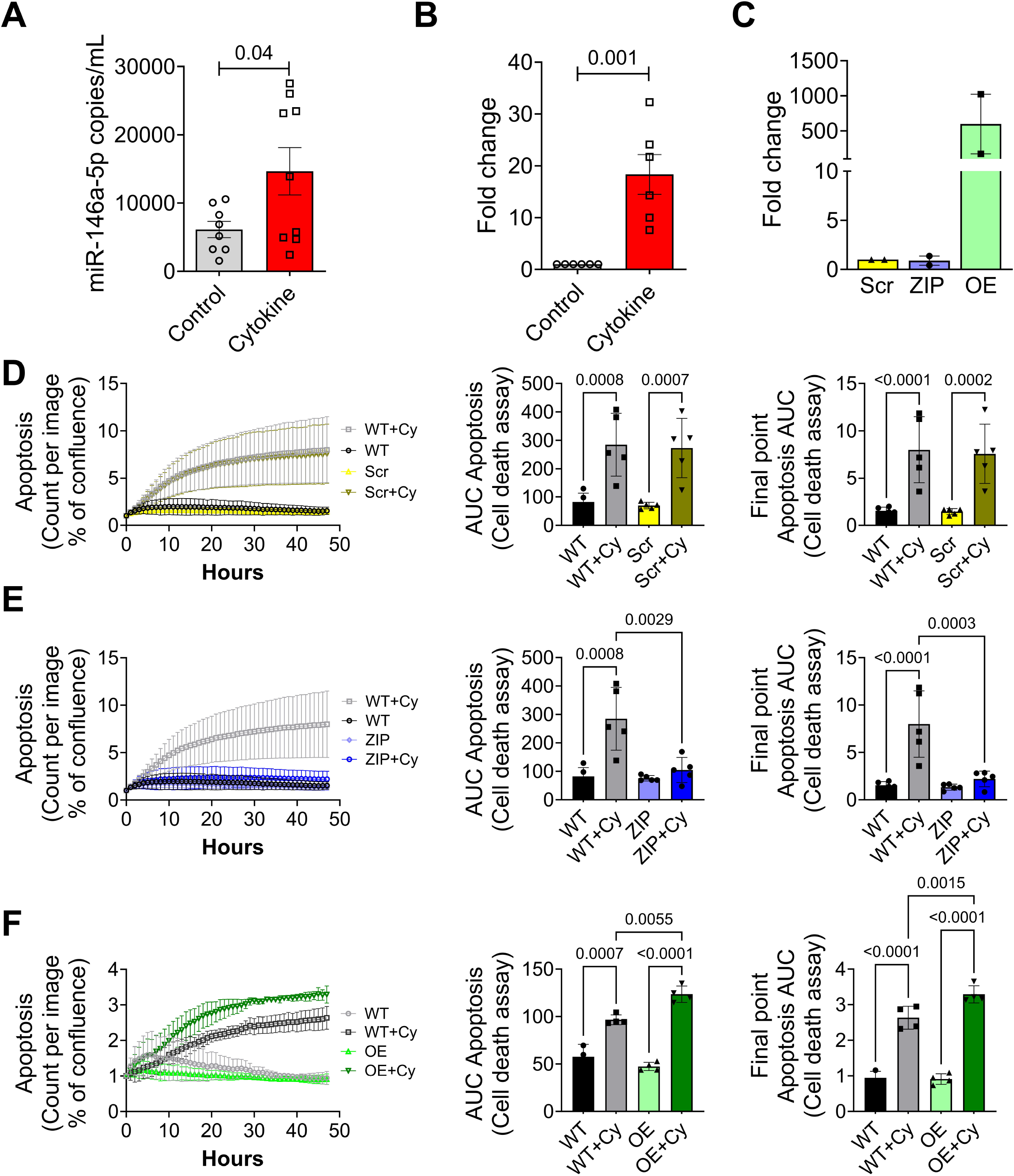
Pro-inflammatory stress increases miR-146a-5p expression levels, leading to increased cell death of MIN6 cells. (A) ddPCR results to determine the expression of miR-146a-5p in mouse islets (n=8). (B) qRT-PCR results to determine miR-146a-5p expression levels in MIN6 cells treated with/without pro-inflammatory cytokines (n=6). (C) qRT-PCR results to determine expression levels of miR-146a-5p in untreated MIN6 stable cells expressing scrambled sequence (Scr), inhibition (ZIP), and overexpression (OE) of miR-146a-5p (n=2). (D-F) Cell death assessment assay (n=5) using Incucyte comparing WT and Scr groups (D), WT and ZIP groups (E), and WT and OE groups (F). Data is presented as means + SEM. Unpaired t-tests were performed for comparisons with two groups. For all comparisons, *p*-values are indicated above.

To determine how modulation of miR-146a-5p levels impacts inflammation-mediated β cell death, we monitored cell death in real-time using a Sartorius Incucyte system. Exposure to pro-inflammatory cytokines increased cell death in wild type MIN6 cells (without any vector; WT) and in Scr cells (Fig. 1D, Fig. S1D). Importantly, inhibition of miR-146a-5p expression (ZIP) reduced cell death in the presence of cytokines (Fig. 1E, Fig. S1E). Conversely, cell death was heightened in OE cells exposed to pro-inflammatory cytokines (Fig. 1F, Fig. S1F), suggesting a direct influence of miR-146a-5p on the regulation of β cell death and survival. Furthermore, we performed immunoblot to quantify the expression of total and cleaved caspase 3, a molecular marker of apoptosis. miR-146a-5p overexpression resulted in a pronounced increase in the cleaved-to-total caspase-3 ratio, whereas this elevation was largely mitigated in cytokine-treated MIN6 ZIP cells (Fig. S2).

### miR-146a-5p regulates pathways related to β cell function and survival

To understand the impact of miR-146a-5p on gene expression, we performed bulk RNA sequencing on Scr, ZIP, and OE MIN6 cells treated with or without cytokines. After filtering for genes with a minimum of ten read counts in total, 17,696 genes were retained for downstream analysis. Principal component analysis (PCA) clustering of all the samples demonstrated distinct separation among the six experimental groups (Fig. 2A). Among all the experimental groups, OE and OE+Cy cells formed distinct clusters separated from the other groups. In contrast, ZIP and ZIP+Cy cells were clustered closer to the control (Scr) group.

**Figure 2.**
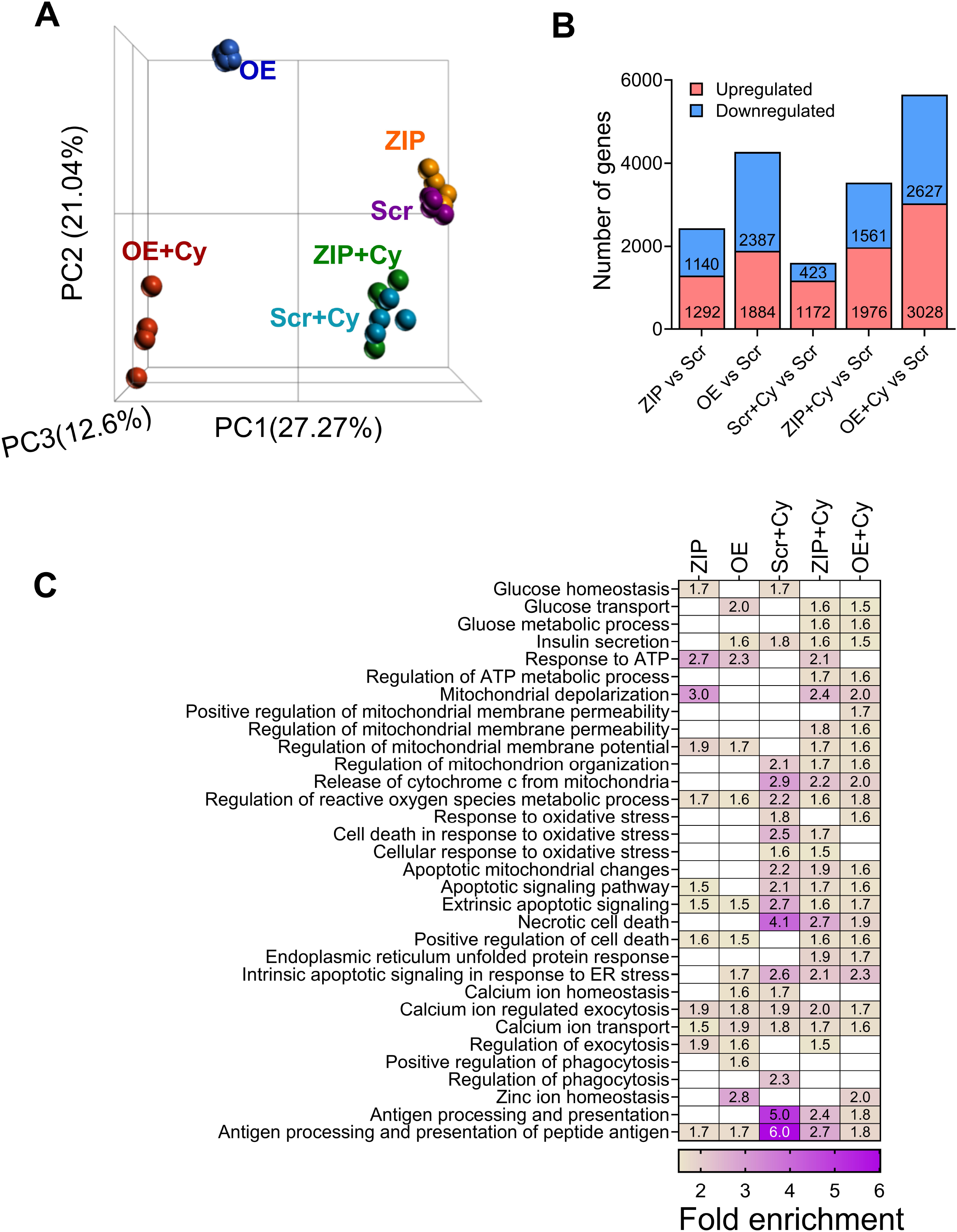
RNA sequencing from MIN6 stable cells show modulation of mitochondrial pathways. (A) PCA plot including all six (Scr, ZIP, OE, Scr+Cy, ZIP+Cy and OE+Cy) comparison groups (n=3-4). (B) Number of differentially expressed genes from each comparison. The comparison groups are indicated below each bar. Red indicates upregulated genes and blue indicates downregulated genes. The numbers included within each bar indicate the number of up and downregulated genes. (C) Functional enrichment results including all comparison groups. Each column represents one group compared to the scrambled (Scr) group. Numbers within each box indicate the fold enrichment value. Higher values are represented in purple and lower values are represented in yellow. Boxes without any numbers indicate no enrichment of the function for that comparison.

First, to assess transcriptomic changes due to miR-146a-5p modulation, we compared ZIP and OE cells to Scr cells and identified 2,432 and 4,271 differentially expressed genes, respectively (Fig. 2B, Supplementary Tables S1a-S1b). Second, to identify genes that are differentially expressed as a result of pro-inflammatory stress, we compared Scr+Cy with Scr and identified 1,595 genes (Fig. 2B, Table S1c). Third, to test if there is a compounding effect of miR-146a-5p expression under inflammatory stress, we compared ZIP+Cy and OE+Cy with Scr as the control group and identified 3,537 and 5,655 genes as differentially expressed, respectively (Fig. 2B, Tables S1d-S1e).

Functional enrichment analysis was performed for differentially expressed genes identified from each comparison group. Applying the filtering criteria for gene ontology terms, we identified terms associated with insulin secretion, glucose, mitochondria, and endoplasmic reticulum, among others. Representative terms and a comparison of their fold enrichment values are provided in Fig. 2C. Notably, enrichment of terms related to mitochondria was observed across all comparison groups. For instance, the term “regulation of mitochondrial membrane potential” was enriched in ZIP vs. Scr and OE vs. Scr both in the presence and absence of cytokines. These findings suggest activation of pathways related to mitochondrial function under pro-inflammatory conditions, with miR-146a-5p potentially playing an independent role in the regulation of mitochondrial function. A complete list of functional terms for all the groups is provided in Supplementary Tables S2a-S2e.

### Knockdown of miR-146a-5p increases insulin secretion

Functional enrichment analysis of the RNA sequencing dataset suggested that miR-146a-5p may be involved in regulating insulin secretion. Therefore, we performed glucose stimulated insulin secretion assays in the MIN6 stable cell lines. In non-cytokine treated cells, we observed a significant increase in insulin secretion under both low and high glucose levels when miR-146a-5p was inhibited compared to Scr cells (Fig. 3A). Under cytokine-treated conditions, we observed a sustained increase in insulin secretion in ZIP cells during high glucose treatment (Fig. 3B). Interestingly, we noted an increase in the insulin content in OE and OE+Cy cells (Fig. 3C); however, insulin secretion was not significantly increased, indicating a potential blockade in the insulin secretory pathway.

**Figure 3.**
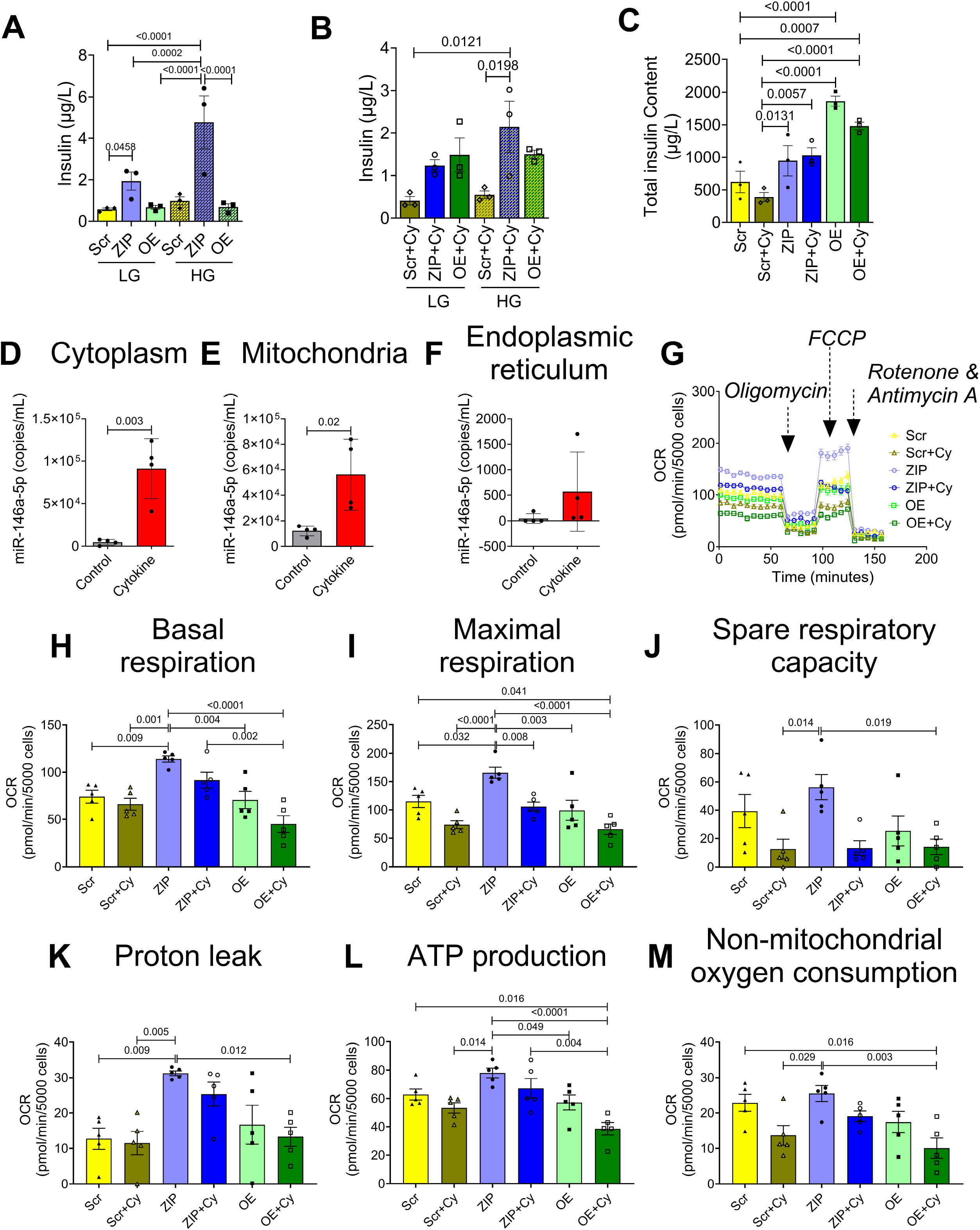
Inhibition of miR-146a-5p increases insulin secretion and respiratory capacity of MIN6 cells. (A) GSIS with low and high glucose in untreated cells. (B) GSIS with low and high glucose in cells treated with proinflammatory cytokines. (C) Total insulin content in treated and untreated cells (n=3). (D-F) ddPCR to determine the expression of miR-146a-5p in cytoplasm (D), mitochondria (E) and endoplasmic reticulum (F) in the presence and absence of pro-inflammatory cytokines (n=4). (G) Normalized oxygen consumption rate calculated over time (minutes) in stable MIN6 cells in the presence and absence of pro-inflammatory cytokines. (H-M) Assessment of other parameters of mitochondrial function including basal respiration (H), maximal respiration (I), spare respiratory capacity (J), proton leak (K), ATP production (L), and non-mitochondrial oxygen consumption (M) (n=5). Data is presented as means + SEM. One-way ANOVAs with Tukey test were used for comparisons with more than two groups. Unpaired t-tests were used for comparing two groups. For all comparisons, *p*-values are indicated above.

### miR-146a-5p exhibits enrichment in mitochondria during inflammatory stress

Next, to test if miR-146a-5p was enriched in the mitochondria of MIN6 cells, we performed subcellular fractionation to isolate mitochondria, cytoplasm, and endoplasmic reticulum (ER). ddPCR analysis from these fractions revealed that cytokine treatment significantly increased the copy number of miR-146a-5p in isolated cytoplasm (Fig. 3D) and mitochondria (Fig. 3E) of WT MIN6 cells. Although we saw a trend toward an increase in miR-146-5p expression in the ER fraction, it was not significant (Fig. 3F). Interestingly, the average number of copies of miR-146-5p in the ER fraction was strikingly lower than the expression seen in the other two fractions. For example, under inflammatory stress, the ER fraction expressed an average of 567 copies per mL whereas the average copies per mL was 56100 and 91125 in the mitochondria and cytoplasm, respectively. Furthermore, the average baseline expression (without inflammatory stress) of miR-146 was highest in mitochondria (12100 copies), followed by the cytoplasm (4583 copies), and the ER fraction (47.5 copies). These findings further support a key role for miR-146-5p in the mitochondria.

### miR-146a-5p induces mitochondrial dysfunction

To assess the impact of miR-146a-5p modulation on mitochondrial function, we conducted several assays to evaluate mitochondrial metabolism, mitochondrial DNA copy number, mitochondrial membrane potential, mitochondrial ultrastructure, and markers of mitochondrial function in MIN6 stable cells.

To detect differences in mitochondrial metabolism, we measured oxygen consumption rates (Fig. 3G) and individual respiratory parameters (Fig. 3H-M) using the Seahorse XF Cell Mito Stress test in untreated and cytokine-treated Scr, ZIP, and OE cells. The Scr group was used as the reference/control group for all comparisons. Our results indicate that ZIP cells exhibited higher basal respiratory rate, maximal respiration rate, and proton leak. Conversely, these parameters were reduced in OE cells upon cytokine treatment. Additionally, ATP production and non-mitochondrial oxygen consumption rates were significantly lower in OE+Cy cells. These findings collectively suggest that inhibition of miR-146a-5p may partially improve mitochondrial respiratory activity.

We then quantified mitochondria DNA copy number in both cytokine-treated and untreated Scr, OE, and ZIP cells by quantifying the copy number of ND1, a mitochondria-encoded gene, relative to nuclear encoded genes *Tbp* and *Hprt*. Our analysis revealed a significant increase in ND1 in ZIP cells compared to Scr cells; in contrast, overexpression of miR-146a-5p did not induce any change in the copy number of ND1 compared to Scr (Fig. 4A).

**Figure 4.**
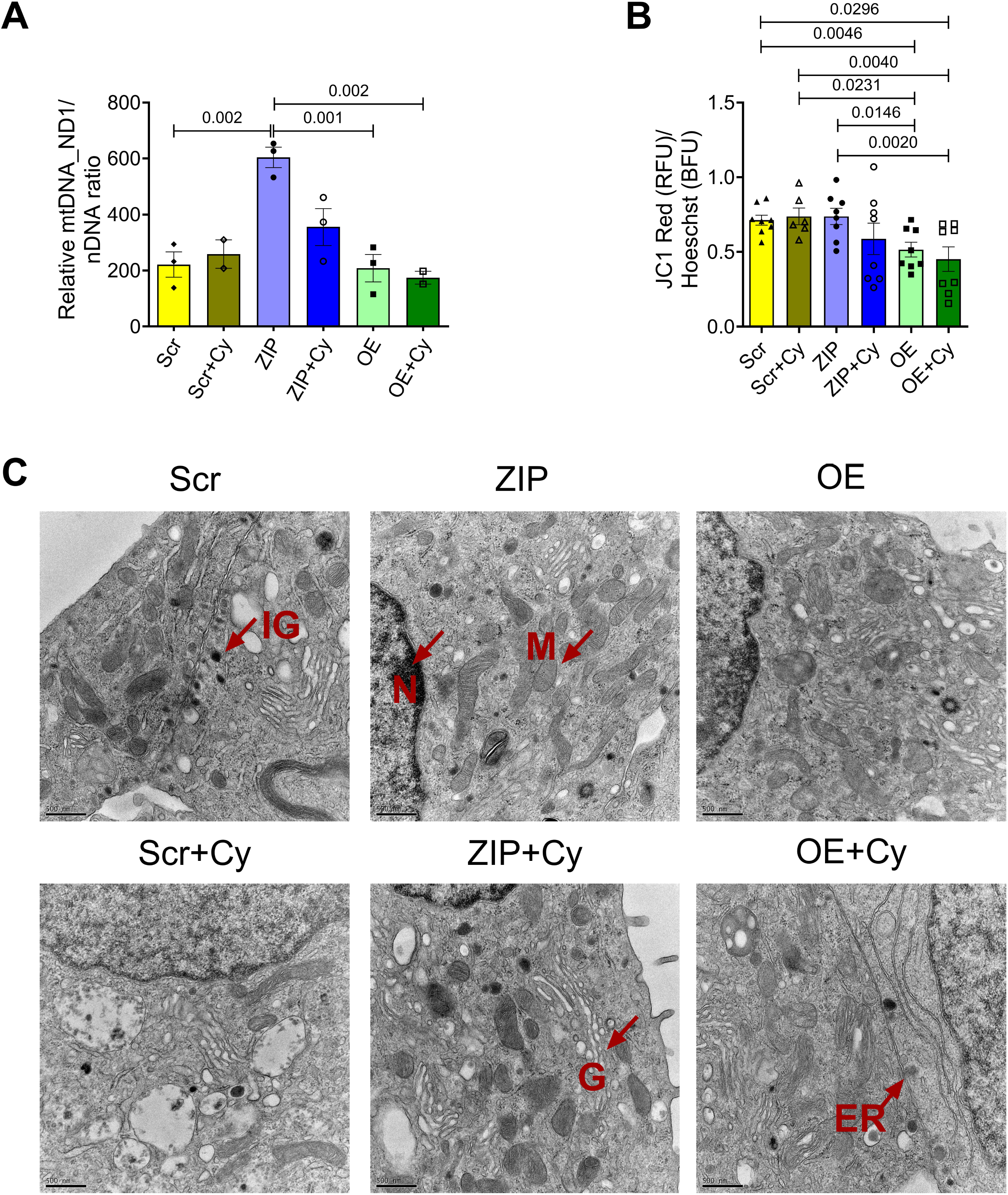
Modulation of miR-146a-5p affects mitochondrial copy number and mitochondrial membrane potential in MIN6 cells. (A) Mitochondria DNA copy number in MIN6 stable cells (n=3) is presented as the ratio of mitochondrial DNA (*Nd1*) to the geometric mean of two nuclear encoded genes (*Tbp, Hprt*). (B) Assessment of mitochondrial membrane potential using JC-1 dye (4 replicates each from two independent experiments) is presented as the ratio of red fluorescence intensity to Hoescht. (C) Electron microscopy images of MIN6 stable cells in the presence and absence of pro-inflammatory cytokines (n=1). M = mitochondria, IG = Insulin granule, N = nucleus, G = Golgi, ER = Endoplasmic reticulum. Scale bar = 500 nm. One-way ANOVAs with Tukey test were used for comparisons with more than two groups. For all comparisons, *p*-values are indicated above.

Using JC-1 dye, we tested whether modulating miR-146a-5p expression levels alters mitochondrial membrane potential (MMP) in MIN6 cells. Because Scr and ZIP cells have GFP reporter, we quantified the red fluorescence intensity alone that could indicate reduced MMP under mitochondrial stress and normalized it to Hoescht. We found that MIN6 cells overexpressing miR-146a-5p exhibited mitochondrial membrane depolarization both at baseline and under inflammatory stress (Fig. 4B). However, ZIP cells did not show a statistical change in mitochondrial membrane depolarization when compared to Scr cells.

Next, to assess whether the overexpression of miR-146a-5p induces ultrastructural changes in β cells, we conducted transmission electron microscopy (TEM) microscopy analysis on the stable MIN6 cell lines. The mitochondrial ultrastructure of ZIP cells remained unchanged by cytokine treatment, and there was an observed increase in the number of mitochondria in both ZIP and ZIP+Cy cells (Figure 4C).

We then examined the expression pattern of an inner mitochondrial protein, TIM23, as a marker for mitochondrial function in our stable MIN6 cell lines. In both the presence and absence of pro-inflammatory cytokines, cells overexpressing miR-146a-5p exhibited significantly lower levels of TIM23 protein compared to Scr and ZIP cells (Fig. 5). This reduction in TIM23 expression, even in the absence of cytokine treatment, suggests that overexpression of miR-146a-5p may drive mitochondrial dysfunction even under basal or unstressed conditions.

**Figure 5.**
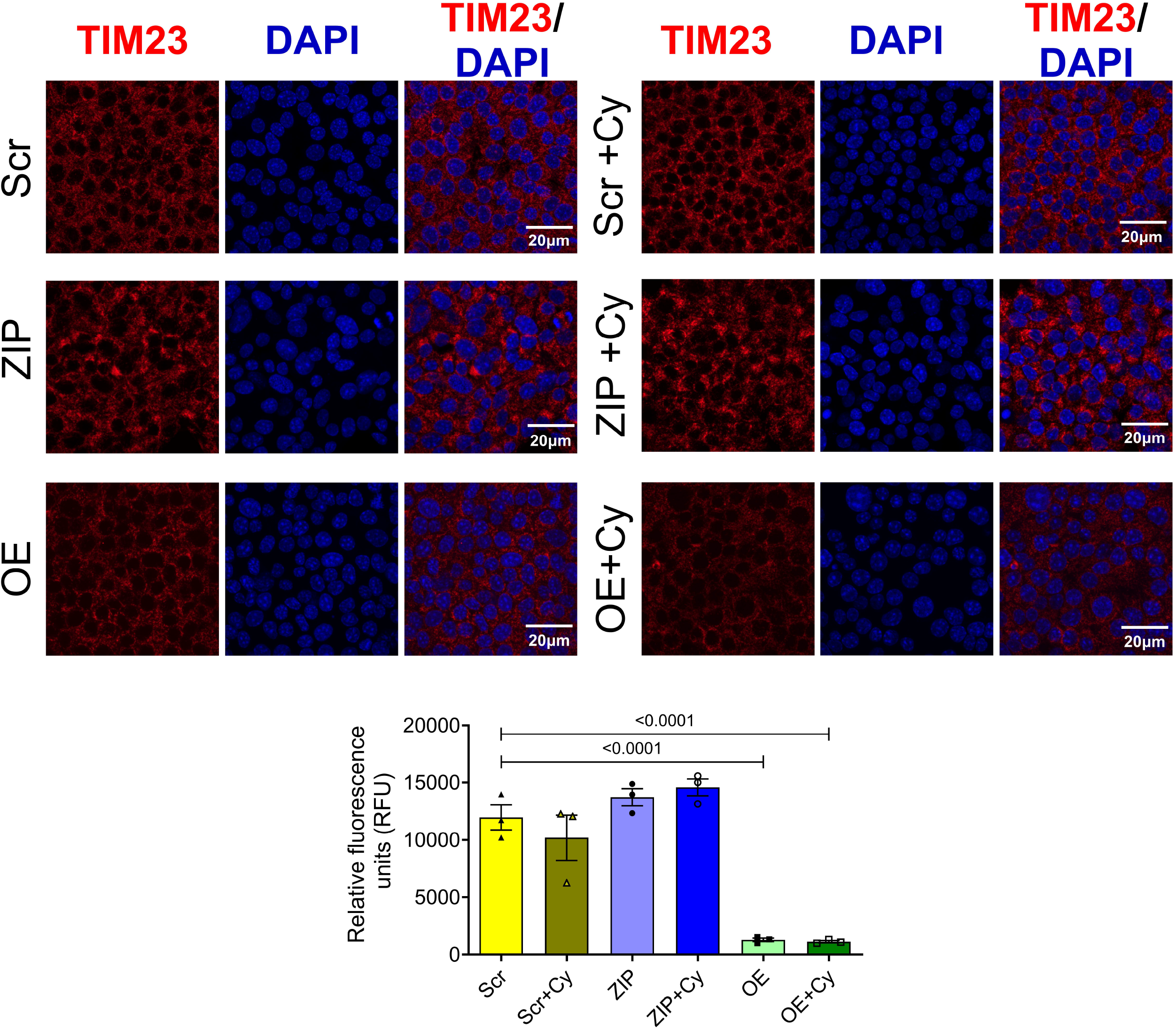
miR-146a-5p overexpression decreases TIM23 expression levels in MIN6 cells. Immunostaining and quantification of TIM23 expression in MIN6 stable cells (n=3). All groups were compared to Scr. Data is presented as means + SEM. Magnification = 40X. One-way ANOVAs with Tukey test were used for comparisons with more than two groups. For all comparisons, *p*-values are indicated above.

### miR-146a-5p targets protein-coding genes related to insulin secretion, apoptosis, and mitochondrial function

Our results collectively indicate that miR-146a-5p is intricately involved in pathways crucial for β cell survival and function, potentially through modulation of mitochondrial function. To identify the gene targets through which these effects may be mediated, we used TargetScan database to predicted 4735 genes as potential targets of miR-146a-5p and overlapped these predictions with our RNAseq dataset, employing several strategies to identify potential targets for validation: 1) We specifically focused on genes included in functional terms (shown in Fig. 2C) containing the words “mitochondria,” “cell death,” “apoptosis,” and “insulin.” This step yielded a total of 109 potential gene targets. 2) To avoid considering genes that were altered only because of cytokine exposure, we considered differentially expressed genes in ZIP vs. Scr and OE vs. Scr comparisons. 3) We only considered gene targets that showed opposite directions of expression in ZIP and OE cells or that were differentially expressed in one group and not in the other group. This strategy reduced the gene list to 22 potential targets. Of these, we chose 9 targets for further validation - *Brsk2, Glrx, Myrip, Rest* (involved in insulin secretion); *Brsk2, Eya1, Sort1, Nrg1, Perp* (involved in apoptosis/cell death), and *Mapt* (involved in mitochondrial membrane potential). Along with these genes, we also included *Gpx1*, a molecule known to be involved in oxidative stress (Fig. 6). qRT-PCR confirmed the differential expression of seven of these ten genes (*Brsk2, Eya1, Gpx1, Mapt, Myrip, Nrg1*, and *Perp*). *Myrip, Gpx1*, and *Nrg1* were upregulated in MIN6 ZIP cells under inflammatory stress, whereas *Brsk2* and *Eya1* were upregulated in cells overexpressing miR-146a-5p. In addition, the change in *Eya1* was significant in OE cells treated with cytokines. *Mapt* was downregulated and *Perp* was upregulated in OE cells treated with cytokines compared with all other groups. Notably, more than 60% of selected target genes were validated, highlighting the potential of miR-146a-5p to modulate pathways involved in β cell function and survival.

**Figure 6.**
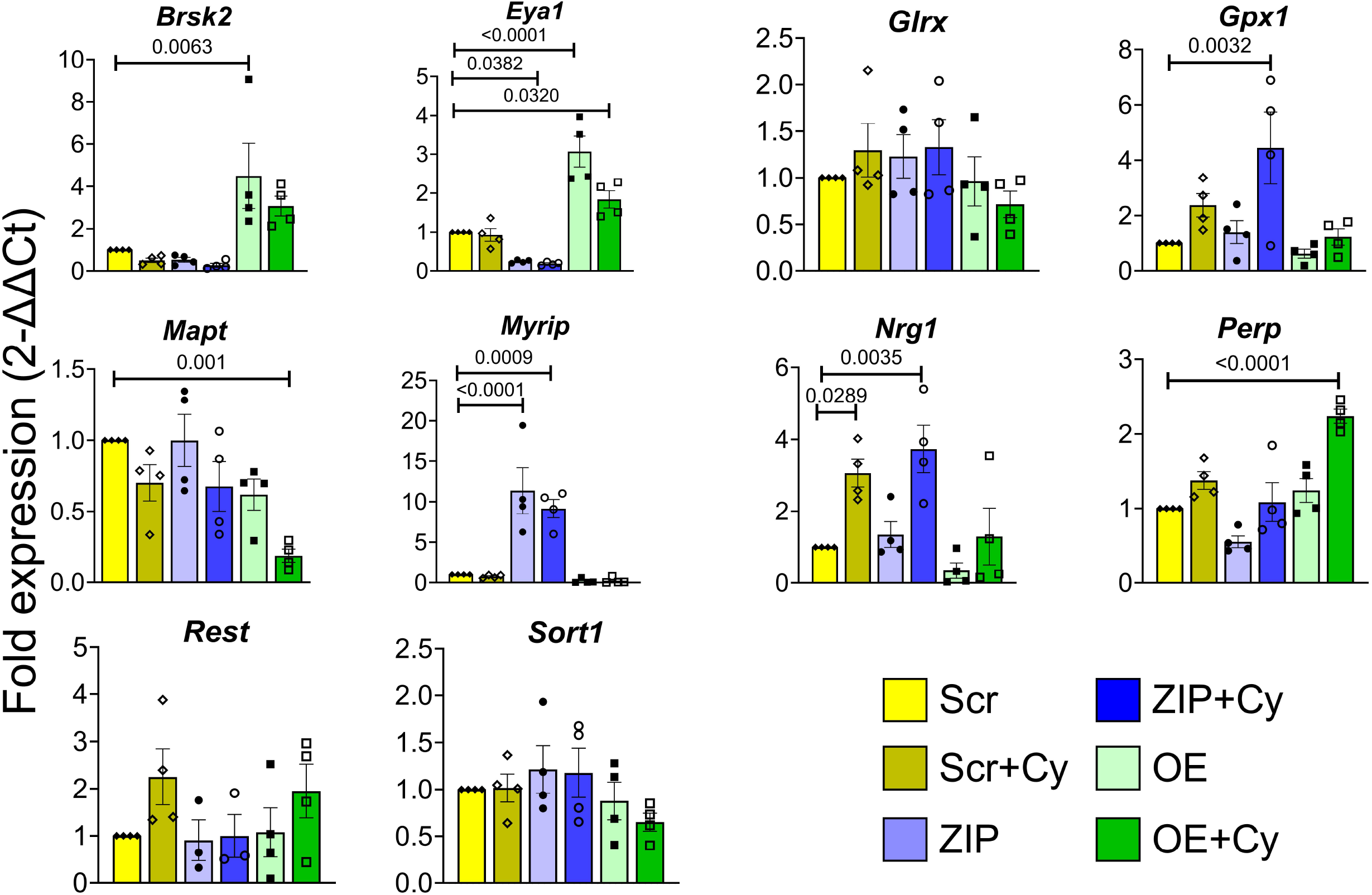
Validation of miR-146a-5p targets. qRT-PCR results of gene targets of miR-146a-5p in MIN6 stable cells (n=4). Data is presented as means + SEM. One-way ANOVAs with Tukey test were used for comparisons with more than two groups. For all comparisons, *p*-values are indicated above.

### Islet mitochondrial dysfunction is observed in mouse models of T1D

To validate the presence of mitochondrial dysfunction in mouse models of T1D during disease progression, we performed immunofluorescence analysis to evaluate the expression of TIM23 and TOM20 (markers of mitochondria function) in islets isolated from nonobese diabetic (NOD) mice at various time points (7, 9, 11, and 13 weeks of age, Fig. 7A). We observed a decreasing trend in TOM20 expression (Fig. 7B) and a significant decrease in TIM23 expression in 13-week-old NOD mice (Fig. 7C), implying that islet mitochondrial function decreases during the progression to T1D. Concurrently, we examined the temporal expression of miR-146a-5p in isolated islets from NOD mice across various time points (5, 7, 9, 11, and 13 weeks of age). Our data demonstrated an elevation in miR-146a-5p expression (Fig. 7D) alongside the decrease in TIM23 and TOM20 expression levels, providing evidence of a correlation between increased miR-146a-5p expression and mitochondrial dysfunction.

**Figure 7.**
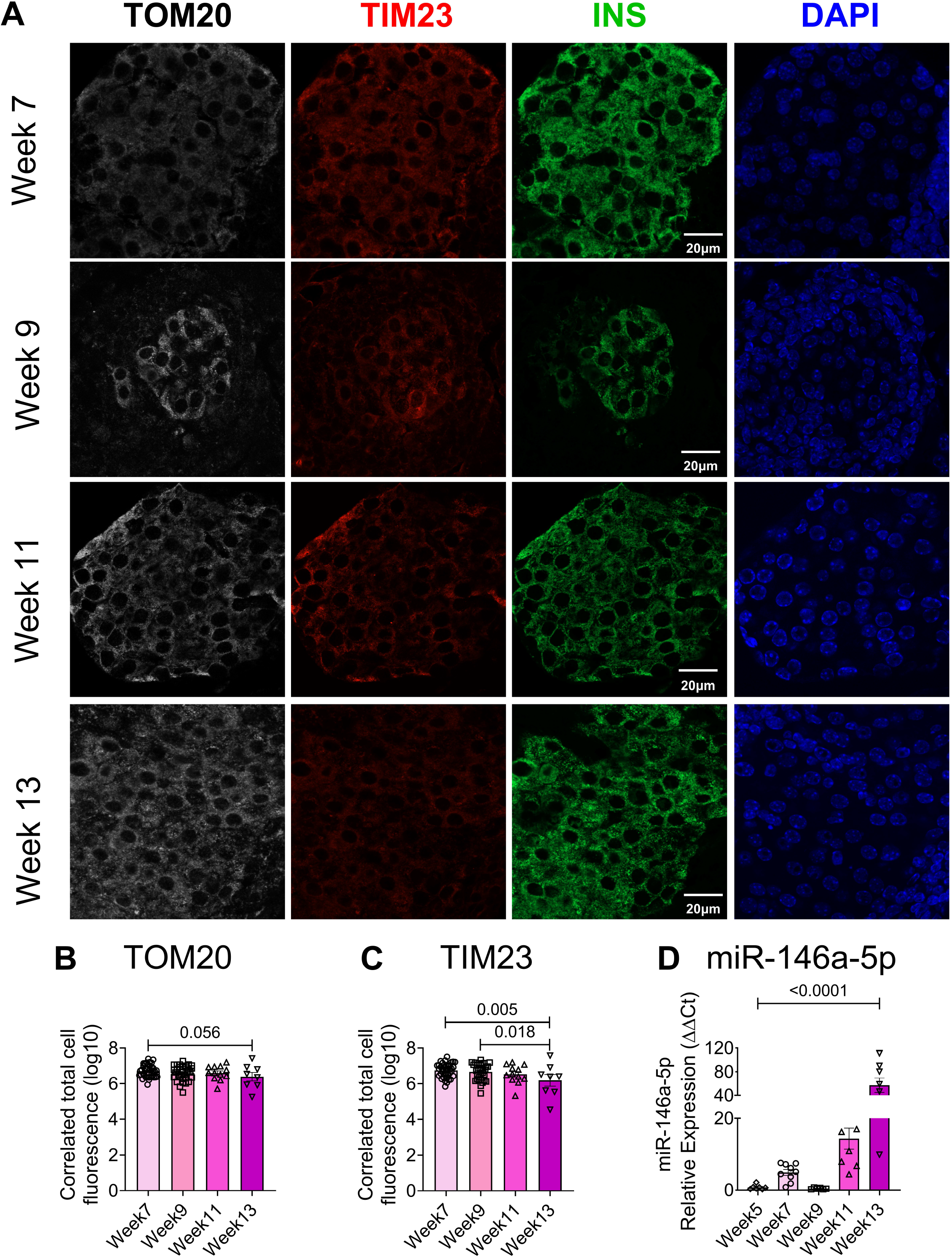
Time-dependent decrease in TOM20 and TIM23 and increase in miR-146a-5p expression in pancreatic islets from NOD mice. (A) Immunostaining of TOM20, TIM23, INS, and DAPI in pancreatic tissue sections of 7-, 9-, 11- and 13-week-old NOD mice (n=8). (B) Quantification of TOM20 and (C) TIM23 using ImageJ. (D) qRT-PCR results to determine time-dependent expression of miR-146a-5p levels in islets isolated from 5-, 7-, 9-, 11- and 13-week old NOD mice (n=8). Magnification = 40X. One-way ANOVAs with Tukey test were used for comparisons with more than two groups. For all comparisons, *p*-values are indicated above.

## DISCUSSION

In this study, we demonstrated that miR-146a-5p is upregulated under conditions of inflammatory stress and that its overexpression leads to mitochondrial dysfunction and apoptosis in β cells, which was reversed by inhibiting miR-146a-5p. These findings represent a novel insight into the role of miR-146a-5p in altering β cell health through the suppression of mitochondrial function.

We have shown previously that miR-146a-5p may serve as a circulating biomarker for T1D (7); however, the molecular mechanisms by which miR-146a-5p contributes to diabetes risk need to be delineated. Roggli *et al*. demonstrated that prolonged exposure of β cells to palmitate and pro-inflammatory cytokines induced alterations in miR-146a-5p expression, including upregulation in human islets, MIN6 cells, and INS1 cells (17). Further, inhibition of miR-146a-5p improved glucose-stimulated insulin secretion and reduced apoptosis in cytokine-treated MIN6 cells. Our findings align with this study by Roggli *et al*., as we observed that inhibition of miR-146a-5p increased glucose stimulated insulin secretion in untreated and cytokine treated cells. We also observed that overexpression of miR-146a-5p in the presence of pro-inflammatory cytokines led to increased apoptosis of MIN6 cells, suggesting that miR-146a-5p may be directly associated with β cell survival.

Functional enrichment analysis of differentially expressed genes identified through RNA sequencing in our MIN6 cell lines provided further insights into the role of miR-146a-5p in pathways associated with β cell survival and function. Notably, modulation of miR-146a-5p resulted in enrichment for terms related to mitochondria, including mitochondrial depolarization, mitochondrial organization, and apoptotic mitochondrial changes. These findings suggest a potential role for miR-146a-5p in T1D pathogenesis, possibly through the modulation of mitochondrial function.

Mitochondria are directly associated with diabetes pathogenesis through alterations in oxidative phosphorylation, ATP production, reactive oxygen species (ROS) formation, and regulation of apoptosis (21–23). This association is supported by several lines of evidence from various studies, including observations of reduced mtDNA copy number in blood samples from individuals with type 2 diabetes (24) and decreased oxidative phosphorylation in the skeletal muscle of insulin-resistant offspring of individuals with T2D (25). Additionally, recent research has highlighted the involvement of miRNAs in mitochondrial function (19). For instance, Dasgupta *et al*. demonstrated the enrichment of miR-146a-5p in mitochondria and predicted mitochondrial genes such as *ND1, ND2, ND4, ND5, ND6*, and *ATP8* as potential targets of miR-146a-5p from 143B cell lines (18). In the context of diabetes, Zhen *et al*. reported a repertoire of mitochondria-associated miRNAs that were dysregulated in mouse liver upon streptozotocin-induced T1D (20). However, the presence and function of mitochondria-associated miRNA in pancreatic islets remain unclear.

To begin to understand if miR-146a-5p expression is detrimental to mitochondrial function, we showed that miR-146a-5p was enriched in mitochondria during inflammatory stress. In line with the observation, seahorse metabolic profiling demonstrated a significant increase in basal respiration, maximal respiration, spare respiratory capacity, proton leak, and ATP production upon inhibiting miR-146a-5p expression. However, these effects were reversed upon overexpressing miR-146a-5p, suggesting its involvement in inducing mitochondrial dysfunction by dampening the activity of the electron transport chain. This hypothesis was well supported by a reduction in mitochondrial membrane potential in cells overexpressing miR-146a-5p compared to Scr cells. This effect was not observed in ZIP cells, suggesting that overexpression of miR-146a-5p may depolarize the membrane, eventually leading to cell death. Since mitochondrial morphology may play a role in metabolic activity (26), we assessed if there were ultrastructural changes in MIN6 stable cells. We observed a potential increase in mitochondria number in ZIP cells, whereas the number of mitochondria were decreased in OE cells treated with/without cytokines. Additionally, OE cells exhibited lower expression of TIM23 both in the presence or absence of cytokines. In a recent investigation, we reported that mitochondrial dysfunction, oxidative stress, and ER-stress pathways were prominently upregulated in prediabetic NOD mouse models, and we identified 14 weeks of age in NOD mice as a critical point for the onset of T1D (27). Consistent with these findings, our current study revealed an inverse correlation between miR-146a-5p and TIM23 expression levels in islets from pre-diabetic female NOD mice across various time points. This finding suggests the potential involvement of miR-146a-5p in mitochondrial dysfunction during T1D progression.

To determine potential mechanisms underlying the impact of miR-146a-5p modulation on β cell function, we validated seven predicted gene targets of miR-146a-5p (*Brsk2, Eya1, Gpx1, Mapt, Myrip, Nrg1*, and *Perp*) via qRT-PCR in our stable MIN6 cell lines. Among these, *Brsk2* (BR Serine/Threonine Kinase 2) is a kinase that plays a key role in insulin secretion and apoptotic responses. In mice, higher expression of *Brsk2* inhibits insulin secretion through interaction with and phosphorylation of PCTAIRE1, while knockdown of *Brsk2* increases serum insulin levels and produces a tendency towards increased β cell size (28). In our experiments, miR-146a-5p overexpression significantly increased *Brsk2* levels. Another identified target of miR-146a-5p, *Myrip* (Myosin VIIA And Rab Interacting Protein), acts as a scaffolding protein to facilitate exocytosis by promoting immediate fusion of insulin granules with the plasma membrane without predocking (29). Higher expression of *Myrip* in our ZIP cells may represent another mechanism through which miR-146a-5p modulates insulin secretion.

*Perp* (P53 Apoptosis Effector Related to PMP22), a protein that acts as an effector in the TP53-dependent apoptotic pathway, has been reported as a mediator of β cell dysfunction and apoptosis, and deletion of *Perp* partially preserves β cell function and mass in glucolipotoxicity settings (30). In our study, *Perp* expression was higher in OE cells than in ZIP cells, and its expression increased significantly under the conditions of cytokine stress, implying a possible mechanism by which miR-146a-5p could mediate β cell apoptosis. *Nrg1* (Neuregulin 1), a member of the epidermal growth factor family, is predicted to be involved in calcium signaling and mitochondrial function. Exposure to recombinant human NRG-1 restores mitochondrial membrane potential, respiration rate, and ATP concentration in cardiomyocytes and skeletal muscle cells in an animal model of heart failure (31). While we observed no change in *Nrg1* expression in OE cells, its upregulation in ZIP cells treated with cytokines suggests that mitochondrial function may be preserved when miR-146a-5p is inhibited.

Together, our results demonstrate a possible new mechanism that defines the significance of miR-146a-5p in the development and progression of T1D.

## EXPERIMENTAL PROCEDURES

### Isolation and treatment of mouse pancreatic islets

C57/BL6J and female non-obese diabetic (NOD) mice were purchased from Jackson Laboratories and maintained at the Indiana University School of Medicine Laboratory Animal Resource Center under pathogen-free conditions. Protocols were approved by the Indiana University Animal Care and Use Committee. To assess the expression of miR-146a-5p during pro-inflammatory stress, pancreatic islets were isolated from 10-12-week-old C57/BL6J mice as described previously (32). After recovering overnight, islets were hand-picked and treated with a cocktail of pro-inflammatory cytokines (33, 34): 5 ng/mL IL1β (Cyt-273-b; ProSpec), 100 ng/mL IFNγ (Cyt-358-b; ProSpec), and 10 ng/mL TNFα (Cyt-252-b; ProSpec). Following 24 hours of incubation with cytokines, islets were washed with PBS and stored at -80 °C. To investigate the temporal expression pattern of miR-146a-5p during the progression to T1D, islets were isolated from female NOD mice at 5, 7, 9, 11, and 13 weeks of age, and miR-146a-5p expression was evaluated. For immunofluorescence analysis, pancreatic tissue sections were collected from female NOD mice at 7, 9, 11, and 13 weeks of age. The sections were then assessed for TOM20 and TIM23 expression using immunofluorescence techniques, following previously established protocols (27).

### Cell lines

MIN6 cells were grown in DMEM (11965-092; GIBCO) containing 4.5 g/L D-glucose, 4 mM L-glutamine, 10% fetal bovine serum (FBS, 35-015-CV; Corning), 100 units/mL penicillin, 100 μg/mL streptomycin (30-002-CI; Corning), and 50 μM β-mercaptoethanol (034461-100; ThermoFisher Scientific). Cells were cultured at 37 °C in a humidified atmosphere containing 5% CO_2_. The medium was changed every other day, and cells were passaged weekly and regularly tested for mycoplasma contamination by PCR-based assay (G238; ABM). To investigate the expression of miR-146a-5p under inflammatory stress, MIN6 cells were cultured in the presence or absence of a combination of three pro-inflammatory cytokines (34): 5 ng/mL IL1β (Cyt-273; Prospec Protein Specialists), 100 ng/mL IFNγ (Cyt-358; Prospec Protein Specialists), and 10 ng/mL TNFα (Cyt-252; Prospec Protein Specialists) for 24 hours.

### Generation of stable miR-146a-5p overexpression or knockdown MIN6 cell lines

MIN6 stable cells with (i) overexpression of miR-146a-5p (OE), (ii) inhibition of miR-146a-5p (ZIP), and (iii) controls with a scrambled sequence bearing no similarity to any known mouse genomic sequence (Scr) were generated using lentiviral vectors. For overexpression of miR-146a-5p, the shMIMIC lentiviral microRNA vector was purchased from Horizon Discovery with Turbo RFP as a reporter (GSH11926-213630770). Lentivector-based scrambled miRZip-000 (MZIP000-PA-1) and miRZip-146a (anti-miRNA; MZIP146a-PA-1) in S1505A-1 construct vector were purchased from System Biosciences. Both Scrambled and miRZIP vectors contained copGFP as a fluorescent reporter. Upon receipt, glycerol *E. coli* stocks for ZIP and Scr were amplified using an LB broth medium with 100 μg/mL carbenicillin for 24 hours in a bacterial shaker at 37 °C. The bacterial culture was pelleted from LB growth medium for all three vectors, and plasmid DNA was isolated using NucleoSpin® Plasmid (NoLid) kit for miniprep (740499.250; Takara Bio) following manufacturer’s instructions. Lentiviruses were generated by transfecting the plasmids using pPACKH1 Packaging Plasmid Mix (LV500A-1; System Biosciences) into a 293T cell line (LV900A-1, 293TN Producer Cell Line; SBI). After 72 hours of culture, viral particles were collected and concentrated using a Lenti-X concentrator (631231; Takara Bio). Lentiviral titers were measured using Lenti-X GoStix (PT5185-2; Clontech, Takara Bio) following instructions provided by the manufacturer. Purified viral particles were transduced into MIN6 cells using TransDux MAX (LV860A-1; System Biosciences) and cultured for 48-72 hours in complete DMEM (11965-092; ThermoFisher Scientific) supplemented with 10% FBS. For clonal selection, transduced cells were cultured with 1 μg/mL puromycin for 7 days. Single clones were picked, seeded in 96-well plates, and cultured with puromycin-containing complete media. The clones were expanded and screened for miR-146a-5p using qRT-PCR, and the clones showing expression patterns of inhibition and overexpression were picked and used for further experiments.

### RNA isolation and droplet digital PCR

Total RNA was isolated from mouse islets, MIN6 cells, and organelle fractions using the miRNeasy mini kit (217004; Qiagen) following the manufacturer’s instructions. RNA concentration was measured using a NanoDrop 2000 (ThermoFisher Scientific). cDNA was synthesized using the TaqMan MicroRNA Reverse Transcription kit (4366596; ThermoFisher Scientific) according to the manufacturer’s instructions.

Droplet digital PCR (ddPCR) was used with a few modifications to measure the expression of mmu-miR-146a-5p, as described previously (39). Briefly, 2 μL of cDNA was added to 10 μL of Bio-Rad ddPCR Supermix for Probes (1863024; Bio-Rad), followed by the addition of 1 μL of TaqMan miR-146a-5p 20X primer mix (4427975; Applied Biosystem); final volume was adjusted to 20 μL using RNase- and DNase-free water. Synthetic miR-146a-5p oligo was used as a positive control, and RNase-free water was used as a negative control. The plates were sealed, and droplets were generated using QX200 AutoDG (Bio-Rad) under RNase-free conditions. PCR was performed using a C1000 Thermal Cycler (Bio-Rad), and droplets were read using a QX200 droplet reader (Bio-Rad). Data was analyzed within Quanta Soft™ Software (Bio-Rad). Data are presented as the number of copies/μL.

### qRT-PCR for miRNA and mRNA

miScript or miRCURY LNA Primer Assay was used to quantify the expression of miR-146a-5p (MS00003535 or YP00204688; Qiagen) in MIN6 cells or mouse pancreatic islets. One μg of RNA was reverse transcribed using the miRScript II kit (218161; Qiagen) or miRCURY LNA RT Kit (339340; Qiagen). PCR was performed using miScript II (218073; Qiagen) or miRCURY LNA (339371; Qiagen) SYBR Green-based PCR kit. At least three replicates were used per condition. PCR reactions were run in duplicates, and thermal cycling conditions were set according to the manufacturer’s instructions. miRScipt II *Rnu*6 (MS00033740; Qiagen) or miRCURY LNA *Rnu6* (YP02119464; Qiagen) was used as the internal reference gene for normalization. Fold-change was calculated using the 2^-ΔΔ^Ct method (35).

To quantify gene expression, 500 ng of total RNA was reverse transcribed using the M-MLV RT kit (28025013; Invitrogen). TaqMan Fast Advanced Master Mix (4444557; ThermoFisher Scientific) and gene-specific probes for *Mapt* (Mm00521988_m1), *Myrip* (Mm00460563_m1), *Brsk2* (Mm01279141_m1), *Glrx* (Mm00728386_s1), *Acvr2b* (Mm00431664_m1), *Rest* (Mm00803268_m1), *Perp* (Mm00480750_m1), *Eya1* (Mm00438796_m1), *Sort1* (Mm00490905_m1), *Gpx1* (Mm00656767_g1) and *Nrg1* (Mm01212130_m1) were used. Thermal cycling conditions specified for the Taqman Fast Advanced master mix were followed. *Actb* (Mm02619580_g1) was used as the internal reference gene for normalization. PCR reactions were run in duplicates, and four replicates per condition were used. Fold-change was calculated using the 2^-ΔΔ^Ct method (35).

### Cell death assessment

Cell death assessment was performed using the Sartorius IncuCyte S3 live-cell analysis equipment (Sartorius, Göttingen, Germany) housed at the Indiana University School of Medicine. Briefly, WT MIN6 cells with no viral vector, and Scr, ZIP, and OE MIN6 cells were plated on 96-well cell culture plates and analyzed starting at ∼40-50% confluency. Cells were transferred to the scanning instrument, and cell media containing dyes for detection of exposed phosphatidylserine, a marker for apoptosis, was added to the cells. Annexin-V green was added to WT and OE cells, and Annexin-V Red was added to WT, Scr, and ZIP cells (1:1000 dilution, #4642 and #4641, Sartorius, Ann Arbor, MI - USA) immediately before time 0 of IncuCyte scan. Cytokines were added or not to the media of each cell line immediately before the first scan. Five images from different areas of each well were collected with a 10X objective every hour for 48 hours, and cell death was quantified as the average number of green or red fluorescent counted objects (excitation: 488nm; emission: 523nm) per unit area at each time point and normalized by percent confluency in each scanned area. Cell death over 48 hours was expressed relative to cell death at time 0 within each replicate.

### mRNA sequencing and data analysis

Total RNA was assessed for quality based on RNA integrity number (RIN) using an Agilent Bioanalyzer 2100. All samples had good-quality RNA, with RIN ranging from 9.6 to 9.9. One hundred nanograms of total RNA was used for library preparation. Briefly, cDNA library preparation included mRNA purification, RNA fragmentation, cDNA synthesis, and ligation of index adaptors. Library amplification was performed following the KAPA mRNA Hyper Prep Kit Technical Data Sheet (KR1352 – v4.17, Roche Corporate). Each resulting indexed library was quantified, and its quality was assessed by Qubit and Agilent Bioanalyzer, and multiple libraries were pooled in equal molarity. The pooled libraries were sequenced with a 2×100 bp paired-end configuration on the Illumina NovaSeq 6000 sequencer using the v1.5 Reagent Kit. Approximately 30M reads per library were generated. A Phred quality score (Q score) was used to measure the quality of sequencing. More than 90% of the sequencing reads reached Q30 (99.9% base call accuracy).

The quality of sequencing data was first assessed using FastQC (Babraham Bioinformatics). Further analysis using Fastq files was performed using Partek Flow 10.0.21.0718. Sequencing files were aligned to the mm10 mouse reference genome using STAR aligner 2.7.3a and mapped to mRNA using RefSeq Transcripts 96 annotation database. Genes with less than 10 read counts in total were removed. Differential expression analysis was performed using DESeq2, and genes with fold-change ≥ 1.5 and *p*-value < 0.05 were considered as differentially expressed. All comparisons were made using Scr as the control group. Raw data along with raw counts and normalized values (median ratio values) are deposited in NCBI GEO under the accession ID GSE255756.

### Functional enrichment and identification of mRNA targets

The DAVID bioinformatics tool was used for functional enrichment analysis (36, 37). Differentially expressed genes from each comparison pair were provided as input for gene ontology analysis. For each comparison, gene ontology terms with fold enrichment value ≥ 1.5 and *p*-value < 0.05 were retained for further analysis. Since the objective of the study was to identify the role of miR-146a-5p in T1D pathogenesis, we focused on gene ontology terms containing the following keywords: mitochondria, endoplasmic reticulum, Golgi, insulin, glucose, apoptosis, cell death, reactive oxygen species, oxidative stress, ATP, zinc, calcium, exocytosis, phagocytosis, and antigen processing and presentation.

To identify the molecular targets that are predicted to bind with the seed sequence of miR-146a-5p, first, *in silico* target prediction was performed with TargetScan 7.2 using a mouse genome as the input (38). We used all the targets, irrespective of the target site conservation status. Targets identified from TargetScan were overlapped with genes included in functional terms containing “cell death,” “apoptosis,” “insulin,” and “mitochondria.” These hits were further filtered to the differentially expressed genes generated from ZIP vs. Scr and OE vs. Scr comparisons. Specifically, we only included genes that showed opposite directions of expression between the two comparison pairs or differentially expressed in one comparison pair but not in the other comparison pair. Three, five, and one representative targets involved in insulin secretion, apoptosis/cell death, and mitochondrial membrane potential, respectively, were selected for validation using qRT-PCR. Although *Gpx1* was not predicted to be a direct target, we included it because of its role in oxidative stress and its opposing expression pattern in ZIP and OE cells.

### Glucose-stimulated insulin secretion

Scr, ZIP, and OE MIN6 cells were plated on 12-well cell culture plates and cultured in DMEM media until 70-80% confluency. The cells were then treated with or without IL-1β, IFN-γ, and TNF-α for 24 hours. Before glucose-stimulated insulin secretion assays were performed, cells were washed twice with freshly prepared glucose-free modified KRBH buffer (5 mM KCl, 120 mM NaCl, 15 mM HEPES, pH 7.4, 24 mM NaHCO3, 1 mM MgCl2, 2 mM CaCl2, and 1 mg/mL radioimmunoassay-grade bovine serum albumin). MIN6 cells were then preincubated for one hour with KRBH containing 2.8 mM glucose. After preincubation, cells were washed twice with glucose-free KRBH and incubated for one hour with KRBH containing 2.8 mM glucose (LG), followed by an additional hour of incubation with KRBH containing 16.7 mM glucose (HG). Supernatant was collected after both 2.8 mM and 16.7 mM glucose stimulations. For total insulin extraction, cells were washed twice with glucose-free KRBH and collected in an alcohol-acidic solution (75% ethanol, 1,5% 1N NaOH, 23.5% deionized water). Secreted insulin and total insulin content were measured using an Insulin ELISA kit (Mercodia; 10-1113-01).

### Isolation of mitochondria, cytoplasm, and ER

Mitochondria and cytoplasm from MIN6 cells were isolated using methods described previously (39). Briefly, 2×10^6^ cells were seeded in a 10 cm culture dish and cultured for 3 days. After 3 days, the cells were replenished with fresh culture media and treated with or without IL1β + IFNγ + TNFa for 24 hours. After the treatment period, the cells were trypsinized with 0.05% trypsin, resuspended with 1X PBS (14190-144; Gibco), and transferred to a 15 mL tube. The cells were pelleted at 215 x g for 5 minutes at 4 °C. The supernatant was discarded, and the cells were resuspended in 500 μL of hypotonic buffer (50 mM HEPES, 1 mM EDTA (pH 8.0), 1 mM Dithiothreitol (DTT), 1X protease inhibitor cocktail (635673; Takara Clontech), 1 mM PMSF, and 1 mg/mL BSA Fraction V) and incubated on ice for 5 minutes. The cells were lysed using digitonin at a final concentration of 0.02% and vortexed every 10 seconds for 5 minutes and mixed with 1:1 (v/v) of 2X isotonic buffer (100 mM HEPES, 2 mM EDTA (pH 8.0), 2 mM DTT, 1X Protease inhibitor cocktail, 2 mM PMSF, 2 mg/mL BSA, and 1.2 M D-Sorbitol) and centrifuged at 700 x g for 10 minutes at 4 ºC. After centrifugation, the supernatant was transferred to a new 1.5 mL tube and centrifuged at 10,000 x g for 15 minutes at 4 ºC. Again, the supernatant was transferred to a new 1.5 mL tube (cytoplasmic fraction) and stored at -80 ºC. 1X isotonic buffer was added on top of the mitochondria pellet and centrifuged at 10,000 x g for 5 minutes at 4 ºC. The supernatant was removed and the pellet containing mitochondria was stored at -80 ºC until further use. To isolate the ER fraction, MIN6 cells were cultured at a density of 3×10^6^ cells in two 10 cm culture dishes for three days. After this initial culture period, the medium was refreshed, and the cells were treated with or without proinflammatory cytokines for 24 hours. Cells were then harvested and subjected to ER isolation according to the manufacturer’s protocol provided with the Minute ER Enrichment Kit (Invent Biotechnologies, Inc., Cat# ER-036). To identify the expression of miR-146 in the organelles, RNA was isolated using the Qiagen miRNeasy kit as described above.

### Mitochondrial metabolic assay

MIN6 stable cell lines were subjected to the Seahorse XF Cell Mito Stress test (103015-100; Agilent). The cells were seeded in the Seahorse XFe 96-well plate at 5,000 cells/well and cultured for 24 hours. The medium was changed the following day, and the cells were treated with pro-inflammatory cytokines for an additional 24 hours. Forty-eight hours post seeding, the growth medium was exchanged for glucose-free, Seahorse Phenol red-free DMEM with 200 mM glutamine, and 100 mM pyruvate, and incubated for 1 hour at 37 °C. Subsequently, cells were incubated for 1 hour with a DMEM containing 2.5 mM glucose in a non-CO_2_ incubator at 37 °C. Oxygen consumption rate (OCR) was measured in 2.5 mM glucose-DMEM as the baseline measurement and following 25 mM glucose stimulus, and mitochondrial inhibitors oligomycin, carbonyl cyanide-4-(trifluoromethoxy) phenylhydrazone (FCCP) and rotenone/antimycin A, respectively, according to the manufacturer’s instructions. Cell count/well was calculated using 10X phase contrast images with segmentation (Incucyte). Counts per image were extrapolated to total counts per well using the surface area of each well (10.326 mm^2^) and image area (2.275 mm^2^), i.e., a multiplication factor of 4.538901. The following assay parameters were assessed in the Mito Stress test: basal respiration, maximal respiration, non-mitochondrial respiration, proton leak, ATP production, and spare respiratory capacity. All reactions were run using six replicates per condition, and the assay was repeated five times. OCR data were normalized using the number of cells/well.

### Assessment of mitochondrial DNA copy number

Scr, ZIP, and OE MIN6 cells were cultured in the presence or absence of pro-inflammatory cytokines for 24 hours. Genomic DNA was extracted using a Qiagen DNeasy kit (69504; Qiagen) according to the manufacturer’s instructions and quantified using a Nanodrop spectrophotometer. Ten nanograms of genomic DNA was added to each qPCR reaction (5 μL of BioRad SYBR green 2X master Mix, 0.4 μL of 10 μM forward primer, 0.4μL of 10 μM reverse primer, 2.2 μL of nuclease-free water, and 2 μL of cDNA at 5 ng/μL). Samples were heated to 98 °C for 5 min, then cycled 40 times at 98 °C for 15 sec followed by 60 °C for 30 sec. Samples were then heated to 95 °C, cooled to 60 °C for 1 min, and then re-heated to 95 °C for melting curve analysis. At least three replicates were used, and the samples were run in technical duplicates. Mitochondria-specific DNA *Nd1* was used for estimating the copy number, and the reference genes were nuclear DNA-specific *Tbp* and *Hprt*.

Mitochondrial DNA copy number was calculated using the formula: ΔCt = Ct(nDNA gene)−Ct(mtDNA gene). Copies of mtDNA = 2 × 2ΔCt. Primer sequences for mitochondria-encoded and nuclear-encoded genes are provided in Table S3.

### Mitochondrial membrane potential

Alterations in mitochondrial membrane potential (MMP) were assessed using the JC-1 mitochondrial membrane potential detection kit (ab113850; Abcam). Briefly, Scr, ZIP, and OE MIN6 cells were plated on black-clear bottom 96-well cell culture plates (CLS3614-100EA; Corning) and cultured in DMEM until 70-80% confluency. Then, cells were treated with or without IL-1β, IFN-γ, and TNF-α for 24 hours. After 22 hours, a portion of the cells that were not treated with cytokines was treated with DMEM containing 1 μM FCCP for two hours as a positive control (ab113850; Abcam). Subsequently, cells were washed twice with low-glucose HBSS (14025092; Gibco) containing 0.2% fatty-acid-free BSA (A7030; Sigma-Aldrich) and incubated with 2 μM JC-1 dye for 30 minutes. Cells were then washed twice with low-glucose HBSS with 0.2% BSA, and JC-1 Red fluorescence was measured using a SpectraMax® iD3 (Molecular Devices) at 535 nm (Ex) and 590 nm (Em) to detect alterations in MMP. JC-1 Red fluorescence intensity was normalized to Hoechst fluorescence intensity, a live-cell nuclei-binding dye (62249; ThermoFisher Scientific).

### Electron microscopy imaging

MIN6 Scr, ZIP, and OE stable cells were seeded in a 6-well plate with a density of 2×10^6^ cells/well and cultured in complete DMEM for 48 hours and were treated with pro-inflammatory cytokines (IL1β, IFNγ and TNFα) for 24 hours. MIN6 stable cells not treated with cytokines were considered as control cells. The cells were washed twice with plain DMEM and fixed with 2.5% glutaraldehyde and 4% paraformaldehyde in 0.1 mol/L sodium cacodylate buffer. The cells were collected by scraping the wells, transferred to a 1.5 mL centrifuge tube, and shipped to the Advanced Electron Microscopy Facility at the University of Chicago for electron microscope analysis. Images were analyzed using ImageJ (40) to estimate the number and assess the overall morphology of the mitochondria.

### Immunofluorescence staining

Formalin-fixed paraffin-embedded (FFPE) pancreatic tissues from NOD mice were sectioned with 5 μm thickness and dewaxed at 70 °C according to the methods described previously (27). Briefly, the slides were deparaffinized with xylene twice for 7 minutes and hydrated with a series of different concentrations of alcohol. Antigen retrieval was performed using citrate buffer (H-3300; Vector Laboratories Inc) for 15 minutes at 100 °C, and the slides in the buffer were cooled to room temperature for 20 minutes and washed with 1X PBS twice for 5 minutes. The slides were blocked with animal free blocker (SP-5030; Vector Laboratories Inc) for 1 hour and incubated with primary antibodies against TOM20 (1:200; 42406S; Cell Signaling), TIM23 (1:200; BD-611222; BD Biosciences), or insulin (IR002; Dako) overnight at 4 °C. Then the slides were washed twice with 1X PBS for 5 minutes and incubated with secondary antibodies: Alexa-Fluor 488 goat anti-guinea pig (A11073; Invitrogen), Alexa-Fluor 568 goat anti-mouse (A11031; Invitrogen), and Alexa-Fluor 647 donkey anti-rabbit (A31573; Invitrogen), and counterstained with DAPI for nucleus.

For immunostaining of MIN6 cells, 75,000 cells were seeded per well in an 8-well chamber slide (80841; Ibidi) and incubated for 24 hours. The following day, cells were treated with or without pro-inflammatory cytokines for another 24 hours. Cells were washed twice with 1X PBS and fixed with 4% paraformaldehyde for 30 minutes at 37 °C. After permeabilizing with 0.1% Triton X-100 in PBS (twice for 5 minutes), the cells were blocked using 0.1% FBS in PBS for 1 hour. The cells were then incubated overnight at 4 °C with primary antibody against TIM23 (1:200; BD-611222; BD Biosciences) diluted in PBS supplemented with 5% FBS. The cells were washed three times with 0.1% Triton X-100 in PBS (5 minutes each wash) and incubated with a secondary goat anti-mouse antibody (Alexa 647; 1:200) for 2 hours at room temperature. After washing the cells three times with 0.1% Triton X-100 in PBS, nuclei were stained with DAPI for 15 minutes at room temperature. Cells were washed twice with 0.1% Triton X-100 in PBS and mounted with Fluorsave reagent (345789-20ML; Millipore Sigma). Z-stack images were acquired using an LSM 800 confocal microscope (Carl Zeiss, Germany). Fluorescence intensity was measured using Image J software (NIH).

### Western blotting

Scr, ZIP, and OE MIN6 cells were cultured in a 12-well plate at a seeding density of 4×10^5^ cells/well. Once the cells reached ∼75-80% confluency, they were treated with pro-inflammatory cytokines for 24 hours. After treatment, the cells were washed with PBS and lysed with lysis buffer (50 mM Tris-HCL, 150 mM NaCl, 0.05% Deoxycholate, 0.1% IGEPAL, 0.1% SDS, 0.2% x Sarcosyl, 5% Glycerol, 1mM DTT, 1 mM EDTA, 2 mM MgCL_2,_ Protease inhibitor (04693132001; Roche, Complete mini-EDTA free), and phosphatase inhibitor (04906837001; Roche, PhosphoStop). Protein concentrations were determined using Lowry’s method. Twenty micrograms of protein per sample was heated for 10 minutes at 70 °C and immediately placed on ice. Proteins were separated using a 4-12% Bis-Tris plus gel (NW04122BOX; Invitrogen) and transferred to a PVDF membrane. Membranes were blocked with Intercept blocking buffer (927-70001; LI-COR) and incubated overnight at 4 °C with primary antibodies against β-tubulin (2128; Cell Signaling), caspase-3 (9662; Cell Signaling), and cleaved caspase-3 (9664; Cell Signaling). Membranes were then washed with 0.05% phosphate buffered saline-Tween (PBS-T) for 20 minutes and incubated with donkey anti-rabbit and donkey anti-goat secondary antibodies (LI-COR) for 2 hours. Images were acquired using an LI-COR image scanner, and protein was quantified using ImageStudio software.

### Statistical analysis

All statistical analyses (except for the analysis of RNA sequencing data) were performed using GraphPad Prism version 9.2. For comparison of means between the two groups, a two-tailed unpaired Student’s *t-*test was used. One-way ANOVA was used for the comparison of means among three or more groups. In all tests, *p* < 0.05 was considered to be statistically significant. Data are presented as mean ± standard error of the mean (SEM). Graphs were generated using GraphPad Prism.

## Author Contributions

P.K, F.S, and C.E.M conceived the study and designed the experiments; P.K, F.S, S.A.W, R.C.S.B, G.C, and C.L performed experiments and analyzed and interpreted the data; C.E.M supervised the study and acquired funds to conduct the study; P.K wrote the draft of the manuscript. All authors reviewed and edited the manuscript.

## Funding and Acknowledgements

This work was supported by NIH grants R01 DK093954, DK127308, U01DK127786, and UC4 DK104166 (to CEM), VA Merit Award I01BX001733 (to CEM), 2-SRA-2019-834-S-B, JDRF 2-SRA-2018-493-A-B, and 3-IND-2022-1235-I-X (to CEM), and gifts from the Sigma Beta Sorority, the Ball Brothers Foundation, and the George and Frances Ball Foundation (to CEM). FS was supported by JDRF awards (3-PDF-20016-199-A-N and JDRF 5-CDA-2022-1176-A-N) and by NIH/NIDDK dkNET (U24DK097771). SAW was supported by an F31 Fellowship from the National Institute of Diabetes and Digestive and Kidney Diseases (NIDDK) award F31DK134168 and a TL1 fellowship from the National Center for Advancing Translational Sciences, Clinical and Translational Sciences Award UL1TR002529. The authors acknowledge the support of the Islet and Physiology Core and the Translation Core of the Indiana Diabetes Research Center (2P30DK097512). The authors also thank Kara Orr and Lata Udari from Islet and Physiology Core for their assistance with islet isolation. We thank Dr. Emily Anderson-Baucum (Indiana University School of Medicine) for her suggestions and assistance in editing the text of this manuscript.

