## Supplemental Figures and Legends for "miR-146a-5p mediates inflammation-induced β cell mitochondrial dysfunction and apoptosis"

### Supplementary Figure Legends

**Figure S1. Pro-inflammatory stress increases miR-146a-5p expression in MIN6 stable cells and overexpression leads to cell death.** (A-C) Fold change values from qRT-PCR experiments (n=6) to determine expression level of miR-146a-5p in Scr and Scr+Cy MIN6 cells (A), ZIP and ZIP+Cy MIN6 cells (B), and OE and OE+Cy MIN6 cells (C). (D-F) Incucyte live cell image system to track cell death (represented by green or red fluorescence) over time. Cell death was compared between WT and Scr (D), WT and ZIP (E), and WT and OE (F) treated with or without cytokines. Scale = 400 µm. Unpaired t-test was used for comparing two groups. For all comparisons, *p*-values are indicated above.

**Figure S2. Overexpression of miR-146a-5p increases cleaved caspase-3 levels under pro-inflammatory stress.** (A) Western blot image showing abundance of caspase-3 and cleaved caspase-3 proteins in MIN6 cells (n=4) treated with/without pro-inflammatory cytokines. (B) Quantification of Western blots. Data is presented as means ± SEM and statistical significance was determined by one way ANOVA. For all comparisons, *p*-values are indicated above.

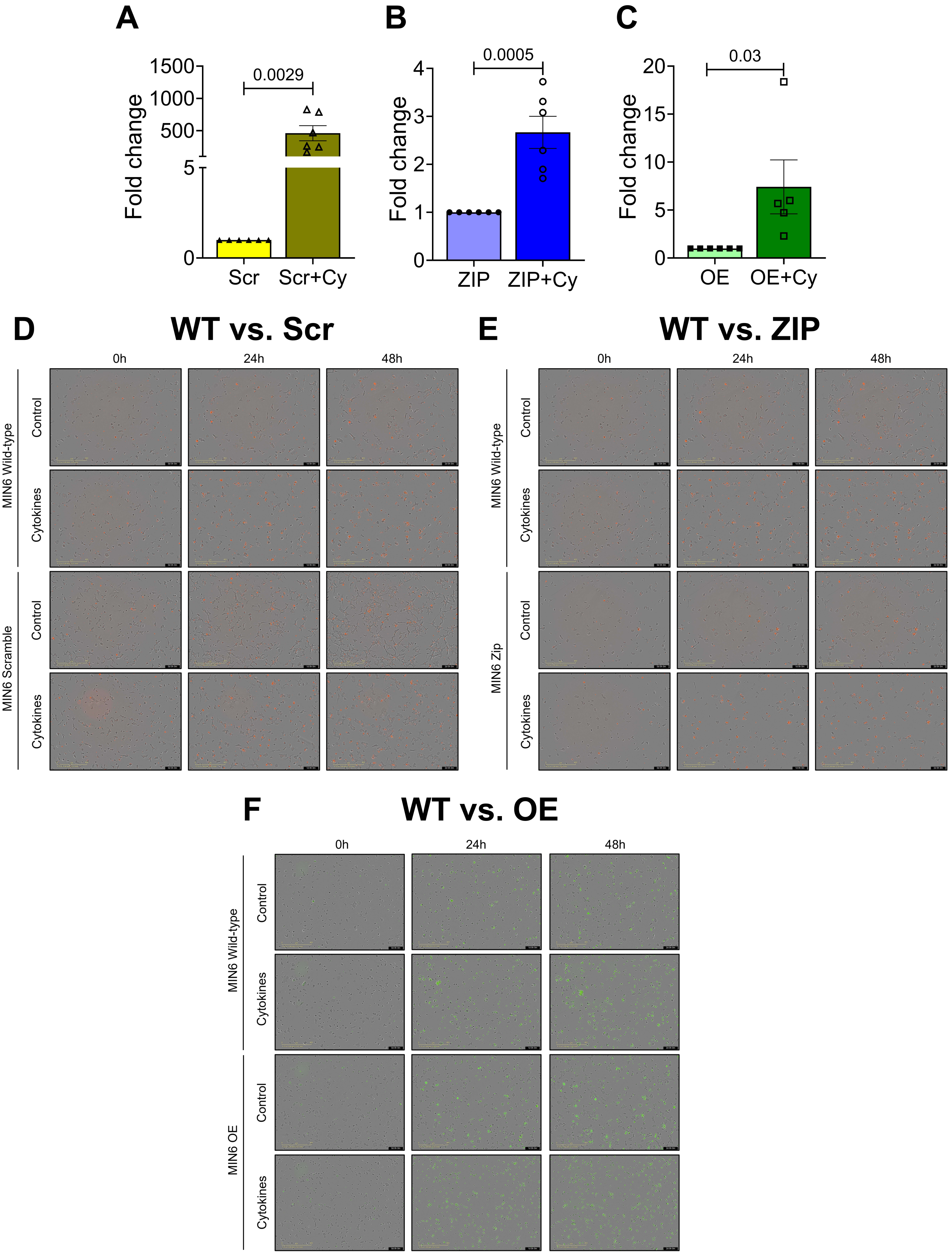
A
Fold change
1500
1000
500
5
0
0.0029
Scr
Scr+Cy
B
Fold change
4
3
2
1
0
0.0005
ZIP
ZIP+Cy
C
Fold change
20
15
10
5
0
0.03
OE
OE+Cy
D
WT vs. Scr
0h
24h
48h
MIN6 Wild-type
Control
Cytokines
MIN6 Scramble
Control
Cytokines
E
WT vs. ZIP
0h
24h
48h
MIN6 Wild-type
Control
Cytokines
MIN6 Zip
Control
Cytokines
F
WT vs. OE
0h
24h
48h
MIN6 Wild-type
Control
Cytokines
MIN6 OE
Control
Cytokines

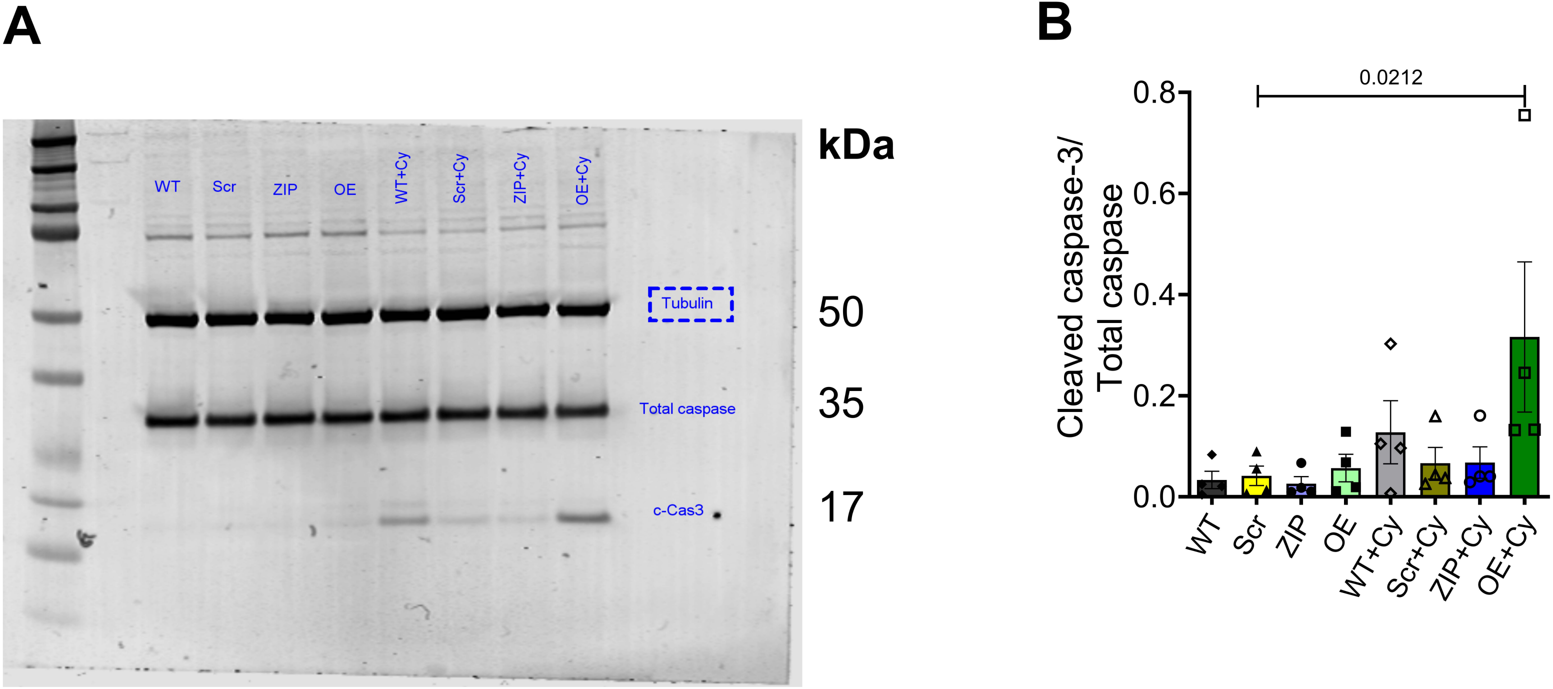
A
WT
Scr
ZIP
OE
WT+Cy
Scr+Cy
ZIP+Cy
OE+Cy
kDa
Tubulin
50
Total caspase
35
c-Cas3
17
B
0.0212
Cleaved caspase-3/
Total caspase
0.8
0.6
0.4
0.2
0.0
WT
Scr
ZIP
OE
WT+Cy
Scr+Cy
ZIP+Cy
OE+Cy
